# Target site and guide RNA multiplexing architecture shape homing gene drive efficiency in *Drosophila suzukii*

**DOI:** 10.64898/2026.07.03.736304

**Authors:** Amarish K. Yadav, Weizhe Chen, Jackson Champer, Maxwell J. Scott

## Abstract

*Drosophila suzukii* (Matsumura, 1931, Diptera: Drosophilidae) is a globally invasive pest of soft-skinned fruits that is currently controlled largely through the use of broad-spectrum insecticides. Increasing resistance to insecticides and regulatory pressures have motivated the development of genetic control strategies. We previously developed a CRISPR/Cas9-based homing gene drive targeting the coding sequence of the female-specific exon of the sex-determination gene *doublesex*, achieving highly efficient inheritance (94–99%) in both male and female germlines. A major limitation of homing gene drives is the formation of resistant alleles that evade cleavage yet retain gene function. Multiplexing guide RNAs (gRNAs) could reduce the formation of such functional resistance alleles. Here, we generated and tested homing constructs expressing one, two, or three gRNAs targeting different regions of the female-specific exon of *doublesex*, including the intron-exon splice junction. A single gRNA targeting the splice junction supported high inheritance in males but showed reduced efficiency in females. Combining this gRNA with a coding sequence-targeting guide further reduced drive efficiency, particularly in the female germline. Constructs expressing two gRNAs performed similarly whether guides were linked by transfer RNA (tRNA) sequences or expressed from independent promoters. Constructs expressing three gRNAs using tRNA processing showed consistently low drive inheritance in both sexes and low frequencies of target-site modification among non-drive progeny, consistent with reduced cleavage activity. Inheritance was significantly higher in male than female germlines for several constructs, indicating that germline context strongly influences drive performance. Our findings show that the strategy used for multi-gRNA expression, target site choice and sex-specific germline environments can influence gene drive efficiency, emphasizing the need to optimize construct design in the target species.

**Author Summary:** Spotted wing drosophila (*Drosophila suzukii*) is an invasive pest that damages soft skinned fruits such as berries and cherries. Control currently relies heavily on insecticides, but resistance and regulatory concerns are increasing the need for alternative approaches. We are developing genetic strategies for suppression of pest populations. We previously developed a gene editing system targeting a female-essential gene that showed very high biased inheritance (gene drive). However, a key challenge is that the system can sometimes create resistant individuals that are unaffected. One possible solution is to use using multiple gRNAs—molecules that direct gene editing to specific DNA sequences—to reduce the formation of these resistant individuals. In this study we found that using more than one gRNA reduced the efficiency of inheritance, especially in females. However, activity could be influenced by exactly where the DNA is targeted and how the guide RNAs are expressed. These results show that gene drive performance depends strongly on biological context and design choices, and that strategies must be carefully optimized in the target species.

## Introduction

Agricultural insect pests cause substantial annual crop losses and threaten global food security, highlighting the urgent need for sustainable, environmentally friendly, responsible and efficient insect pest control strategies (1–3). Genetics-based control approaches provide species-specific alternatives to chemical insecticides and offer the potential for durable population suppression or modification (4–6). Genetic pest control entails releasing of genetically engineered versions of a targeted species with the goal to suppress or modify their wild population (7–9). Significant progress has been made in developing and evaluating genetics-based technologies such as <u>f</u>emale-<u>s</u>pecific Release of Insects carrying a Dominant Lethal (fsRIDL), <u>p</u>recision-<u>g</u>uided Sterile Insect Techniques (pgSIT), X-shredder-based sex-ratio distortion, the toxic male technique (TMT), and CRISPR-based homing gene drives (10–19).

Among these approaches, homing gene drives enable the rapid spread of engineered genetic elements through target populations and therefore require relatively small release sizes compared to conventional genetic control strategies (20). Inspired by naturally present selfish genetic elements, multiple synthetic homing gene drives have been developed and evaluated, including CRISPR/Cas9-based platforms (18, 21–25). In a CRISPR/Cas9-based homing gene drive, germline-expressed Cas9 and constitutively expressed single guide RNAs (sgRNAs) produce double strand breaks (DSB) at genomic target sites. The drive cassette is copied into the homologous chromosome through homology directed repair (HDR), converting germ cells into functional homozygotes and consequently leading to super-Mendelian inheritance of the drive (26, 27). Fluorescent marker genes such as DsRed are commonly incorporated into the drive cassette to facilitate phenotypic screening and strain maintenance (23).

For population suppression, conserved sex determination pathway genes such as *doublesex* (*dsx*) and *transformer* (*tra*) are attractive targets (6, 28). Alternative splicing generates sex-specific isoforms, enabling selective disruption of female development while minimizing impacts on male viability (29, 30). Homing gene drives targeting *dsx* and *tra* have been developed in multiple pest species, including the Mediterranean fruit fly *Ceratitis capitata* (31) and *Drosophila suzukii* (23). *Drosophila suzukii* (commonly referred to as spotted wing drosophila or SWD) is a highly destructive invasive pest of soft-skinned fruits, including cherries and berries. Unlike most drosophilid species, *D. suzukii* females possess a serrated ovipositor that enables oviposition into ripening fruit, where larval development causes direct economic damage and crop loss (32). Current management strategies rely heavily on chemical insecticides; however, the rapid emergence of insecticide resistance, combined with environmental and regulatory concerns, underscores the urgent need for alternative control approaches (33–35). In addition to homing gene drive, fsRIDL (36) and pgSIT (37) strains have been developed and evaluated for this species.

We previously reported a CRISPR/Cas9-based split homing gene drive targeting the female-specific exon of *dsx* in *D. suzukii* (23). In a split homing gene drive, Cas9 and sgRNA components are unlinked (generally on separate chromosomes), in contrast to autonomous or full drives where drive components are genetically linked (25, 38, 39). This facilitates independent optimization of the Cas9 and sgRNA components and provides an important safeguard for laboratory experiments as spread of the drive following an accidental release would be limited (38, 40). The *dsx* homing gene drive demonstrated strong inheritance bias and population suppression potential but relied on a single sgRNA target site (23). With a single sgRNA, the potential for drive resistant allele development is a major concern (41). The DSB repair through error-prone pathways such as non-homologous end joining (NHEJ) can introduce small insertions or deletions that disrupt sgRNA recognition while preserving gene function. These functional resistant (r1) alleles can accumulate over time and impede the drive propagation (42, 43). One approach to reduce the likelihood of r1 alleles accumulating is to target functionally conserved regions where small changes in the encoded amino acid sequence are not tolerated (18). The region we targeted in *D. suzukii dsx* does appear to be functionally constrained as a mutant with an in-frame 3 bp deletion at the target site behaved as a null allele (23). However, it is possible that r1 alleles could still arise such as by small in-frame insertions or a SNP in the seed region or PAM sequence. Another approach is to use multiple sgRNAs to avoid the accumulation of ‘r1’ alleles as this would require resistance to cleavage at each target site simultaneously (44, 45). In addition, simultaneous cutting at multiple sites followed by repair through non-homologous end joining (NHEJ) is expected to generate large deletions that are more likely to disrupt gene function and be rapidly eliminated from the population due to strong fitness costs. Multiple sgRNAs can be expressed either using separate RNA polymerase III promoters for each sgRNA (46) or from a single polycistronic transcript in which individual sgRNAs are separated by tRNA or ribozyme sequences (47). Processing of this transcript by the endogenous tRNA processing machinery or self-processing ribozymes releases functional individual sgRNAs.

The aim of this study was to evaluate homing gene drive constructs expressing two or three sgRNAs targeting *dsx* in *D. suzukii*. These sgRNA were multiplexed using tRNA-based processing systems or expressed from independent *U6* gene promoters. In addition, we also established a single sgRNA homing construct, which targets the functionally constrained *dsx* female splice junction (intron exon boundary), rather than the central coding region targeted previously. Multiplexed homing strategies have reduced resistance allele formation in *D. melanogaster* (44, 48) and mosquitoes (41, 46, 49), and here we extend these approaches to a major agricultural pest species to evaluate drive performance, resistance dynamics, and population suppression potential.

## Results

### Design of homing constructs with multiplexed-sgRNAs targeting *doublesex* (*dsx*)

A homing suppression drive targeting the central coding region of the female-specific exon of *doublesex* (*dsx*) with a single sgRNA was previously established in *D. suzukii* (23). A multiplexed-sgRNAs homing drive targeting the *dsx* splice junction in *D. melanogaster* showed minimal accumulation of functional resistance alleles (24). Guided by these studies, here we designed and developed a variety of homing constructs expressing single and multiple sgRNAs targeting the intron-exon junction and coding sequence (CDS) of the *dsx* female-specific exon in *D. suzukii* (Fig 1A).

**Fig 1.**
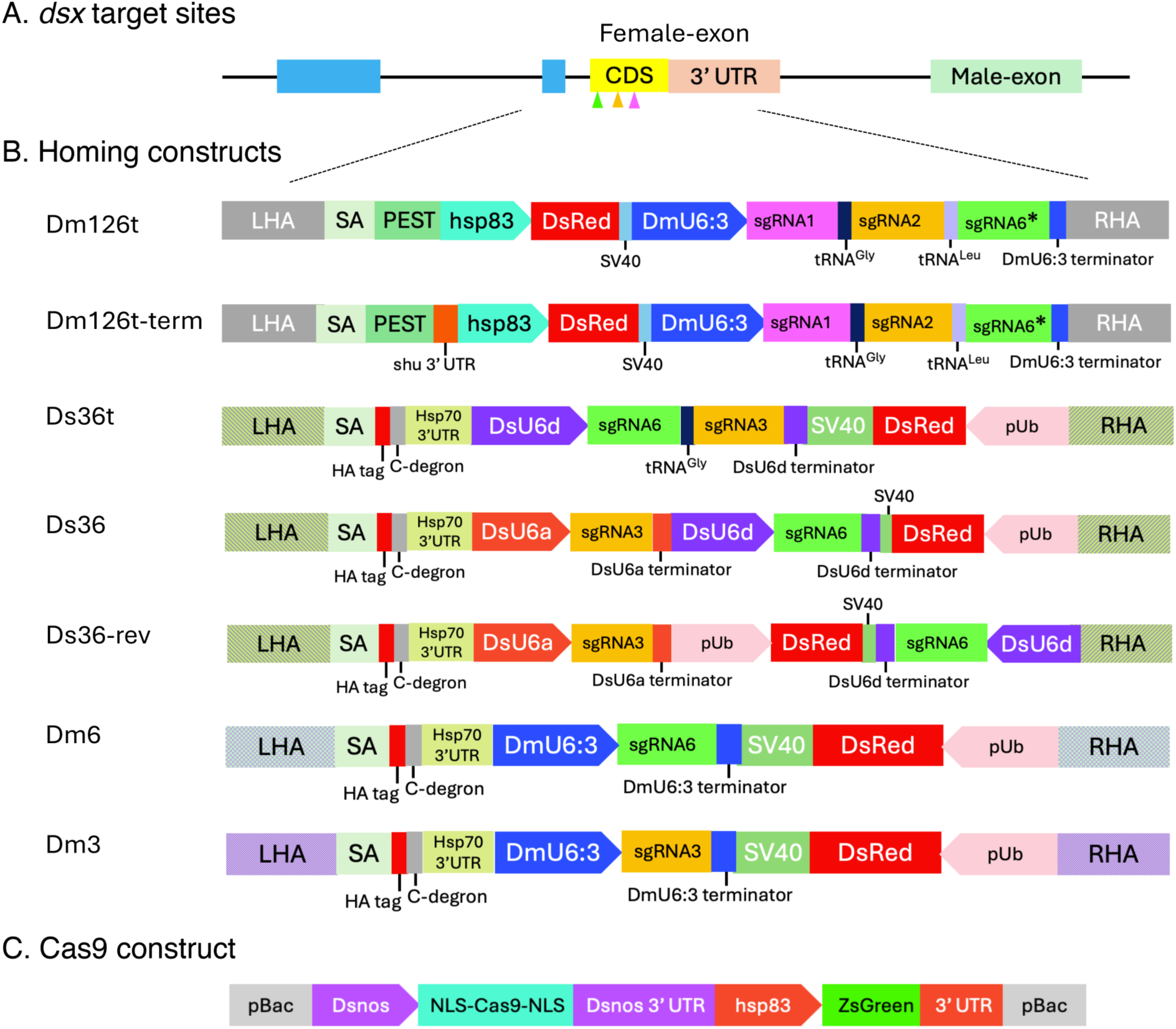
Design of split homing gene drive constructs targeting the *doublesex* (*dsx*) locus in *Drosophila suzukii*. (A) Schematic representation of the *dsx* gene structure and sgRNA target sites (not to scale). Blue boxes indicate exons common to both sexes, while the female-specific exon (coding sequence and 3′ untranslated region) and male-specific exon are shown separately. Colored arrowheads denote sgRNA target sites within the female-specific exon. (B) Structure of homing gene drive constructs and corresponding transgenic strains. Constructs Dm126t and Dm126t-term express three sgRNAs multiplexed using tRNA sequences under the *D. melanogaster U6:3* promoter. Ds36t expresses two sgRNAs linked by a tRNA under the *D. suzukii U6d* promoter. Ds36 and Ds36-rev express two sgRNAs from independent *D. suzukii* U6 promoters arranged either back-to-back (Ds36) or separated (Ds36-rev). Dm6 and Dm3 express a single sgRNA under the *D. melanogaster* U6:3 promoter. Left and right homology arms (LHA and RHA) flank the constructs and differ slightly among designs, reflecting the positions of the outermost sgRNA cut sites. Additional elements include splice acceptor (SA), degradation sequences (e.g., PEST or C-degron), promoter and terminator elements, and a DsRed fluorescent marker for screening. (C) Germline Cas9 expression construct. Cas9-2xNLS is expressed under control of the *D. suzukii nanos* promoter and 3′ untranslated region, and fused to nuclear localization signals to ensure efficient nuclear targeting.

Constructs Dm126t and Dm126t-term were designed to express three sgRNAs (sgRNA1, sgRNA2, and sgRNA6*) multiplexed using a tRNA processing system and driven by a *D. melanogaster U6:3* promoter-terminator (Fig 1B). These sgRNAs target the splice-junction (sgRNA6*), middle (sgRNA2) and downstream CDS (sgRNA1) of the female-specific exon (S1 Fig 1A). Distinct guide-RNA scaffold sequences were used to minimize repetitive sequences within the constructs (S1 Fig. panel B). In both constructs, sgRNA1 was positioned immediately following the *U6:3* promoter, while the sgRNA2 and sgRNA6* were separated using tRNA^Gly^ and tRNA^Leu^ sequences, respectively. To avoid dominant female-sterility, an efficient splice acceptor site from a *D. melanogaster myosin* gene was used as previously in *D. suzukii* and *D. melanogaster* (23, 24). The constructs Dm126t and Dm126t-term differ only in their myosin splice-acceptor site structure located downstream of the left homology arm. To minimize inhibition of the DSX protein produced by the wild type *dsx* gene, both constructs contain an in-frame PEST sequence to destabilize the truncated DSX protein produced by the drive allele (24). Dm126t lacks a 3’ UTR with a known polyadenylation site, whereas Dm126t-term includes a 3’UTR from the *D. melanogaster shu* gene that contains predicted polyadenylation sites (Fig 1B). Both constructs contain a *hsp83* promoter driven DsRed fluorescent marker gene. Following microinjections of homing constructs and Cas9 ribonucleoprotein complexes targeting the outermost cut sites into precellular *D. suzukii* embryos, transgenic lines were established (S1 Table).

Construct Ds36t was designed to express two sgRNAs (sgRNA6 and sgRNA3) under the *D. suzukii U6d* promoter-terminator and multiplex using a tRNA^Gly^ sequence (Fig. 1 B). These sgRNAs target sites about 50 bp apart, with sgRNA6 targeting the splice-junction and sgRNA3 targeting the central coding region (S1 Fig. A). sgRNA6 targets an identical sequence as sgRNA6* but has a different scaffold, and sgRNA3 largely overlaps with sgRNA2 (S1 Fig.). The sgRNA6 is designed to disrupt a highly conserved intron-exon junction, which could minimize functional resistant alleles generation (18). Distinct scaffold sequences were used to avoid repetitive DNA sequences (S1 Fig. B). From seven independent lines, one line, was established and used for homing assessment (S1 Table).

Two additional constructs, Ds36 and Ds36-rev, were designed to express sgRNA3 and sgRNA6 using independent *D. suzukii U6a* and *U6d* promoter-terminators (Fig. 1B). Unlike the constructs described above, the guide-RNAs scaffold sequences are identical (S1 Fig. 1). In Ds36, the *DsU6a* and *DsU6d* promoters are in a tandem orientation, whereas in Ds36-rev these promoters are spatially separated and in opposite orientations (Fig 1B). Single strains from four Ds36 lines and five Ds36-rev lines were selected for drive assessment (S1 Data).

Finally, for comparison with the single sgRNA3 *dsx^SA^*drive evaluated previously (now renamed Dm3) (23), Dm6 was designed to express only sgRNA6 under the *D. melanogaster U6:3* promoter (Fig 1B). A strain was established from two independent lines (S1 Data). The Ds36t, Ds36, Ds36-rev and Dm6 constructs all contain the myosin splice acceptor sequence and a polyubiquitin promoter-driven DsRed marker gene (Fig. 1B).

Correct site-specific insertion of all homing constructs into the *dsx* locus was confirmed by PCR and sequencing (S2 Fig). As expected, hemizygous males and females were fertile, whereas homozygous male and female were sterile and exhibited intersex phenotypes and malformed genitalia. Homozygous male sterility is likely due to disruption of the male-mode of *dsx* RNA splicing due to the use of myosin splice acceptor site (23).

### Assessment of split homing gene drive efficiency

Homing efficiency was evaluated using a split homing gene drive approach as described in previous studies (23, 25). Two previously characterized Cas9 strains, X-linked (#27A1) and autosomal (#25A2), were used as Cas9 sources (Fig. 1C). Both lines express NLS-Cas9-NLS in germ-cells under the *D. suzukii nanos* gene promoter and 3’UTR and have comparable level of Cas9 activity (23). Hemizygous males of the homing strain were crossed with homozygous Cas9 virgin females to obtain ‘male-drive’ and ‘female-drive’ progeny carrying both the sgRNA and Cas9 transgenes. Male-drive and female-drive were subsequently crossed with wild-type flies to measure homing efficiency by scoring the inheritance of the DsRed marker. Successful homing was defined as inheritance exceeding the expected Mendelian frequency of 50% (7, 50).

As there were no significant differences in drive efficiency using the X-linked or autosomal Cas9 lines, in this section we report only the results obtained with the X-linked Cas9 line, but full results are provided in Supplementary (S2 Data). Except for female drive for the Dm126t line, all lines produced significant gene drive (i.e. >50% inheritance) (Fig. 2, S3 Data).

**Fig 2.**
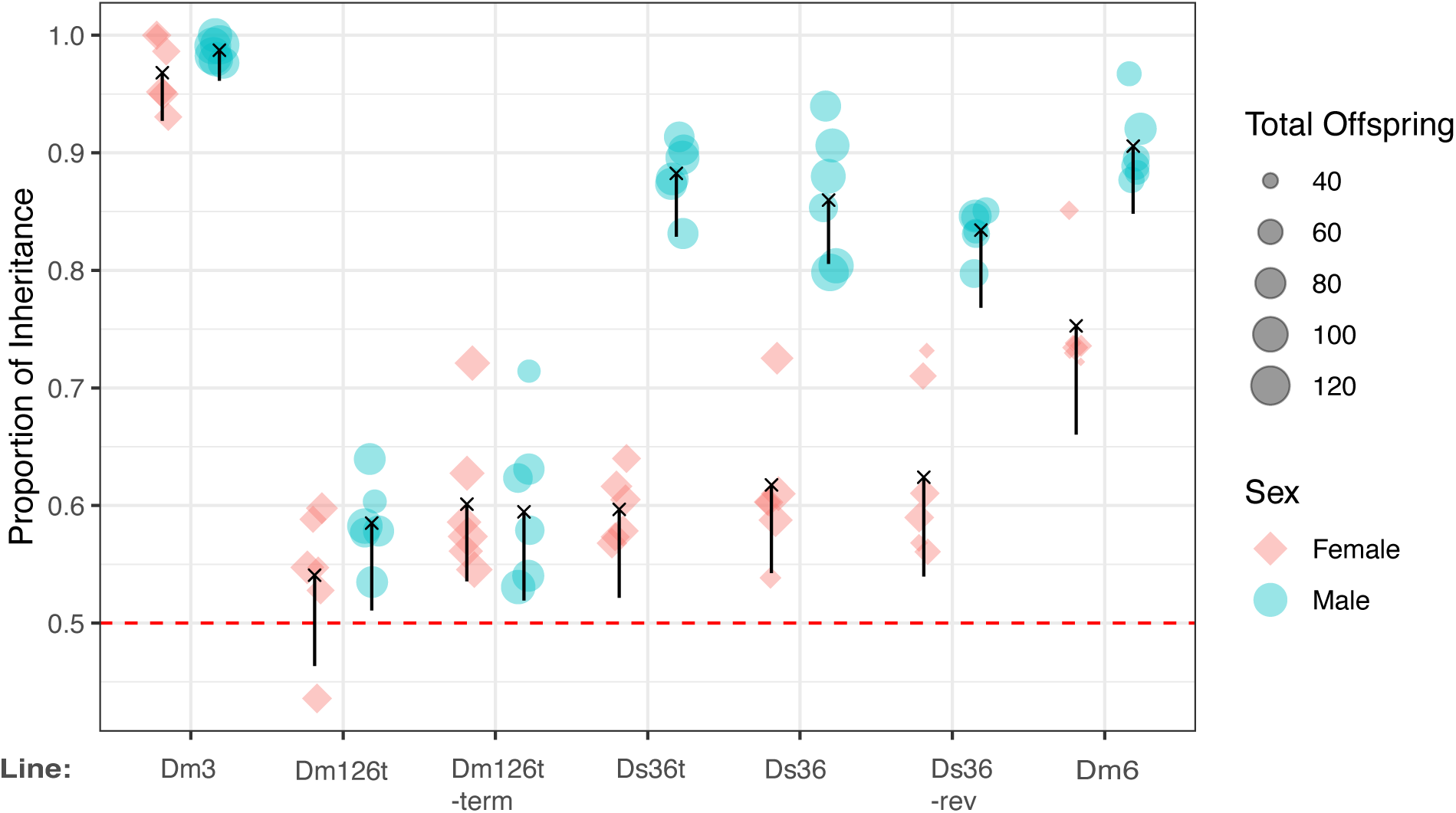
Homing gene drive inheritance varies with sgRNA design and differs between male and female germlines. Homing gene drive inheritance for constructs targeting the *doublesex* locus in *Drosophila suzukii*. Each point represents a single replicate cross, with point size proportional to the total number of progeny scored. Male-drive (teal circles) and female-drive (pink diamonds) crosses are shown separately for each line. Black crosses indicate mean inheritance, with vertical bars representing adjusted one-sided Bonferroni intervals. The dashed red line indicates the expected Mendelian inheritance frequency (50%). The gene drive inheritance data inheritance for the Dm3 line (previously called 23AI1) that expresses sgRNA3 alone is from our previous study (23).

The three-gRNA strains Dm126t and Dm126-term, produced drive inheritance frequencies of 58.7% and 60.5% in male-drives and 54.1% and 60.2% in female-drives, respectively (Fig. 2). There was no significant difference in male and female-drive for each strain. For the two-sgRNA tRNA multiplexed strain Ds36t, male-drive was significantly higher with observed inheritance of 88.2% (Fig 2, S3 Data, *p*<0.001). However, female-drive was comparable to Dm126t and Dm126-term with 60.1% inheritance. For the two-*U6* promoter constructs, the Ds36 line that has tandem *U6* promoters, exhibited drive inheritance of 86.4% (male-drive) and 61.1% (female-drive) using X-linked Cas9, with similar values for the autosomal Cas9 (S2 Data). For the Ds36-rev line where the *U6* promoters are spatially separated, inheritance rates of 83.4% for male-drives and 62.9% for female-drives were observed (Fig 2 A-B). For both lines, drive efficiency was significantly higher in males than females (Fig 2, S3 Data, p<0.001). The two gRNA lines, Ds36t, Ds36 and Ds36-rev were comparable for both male and female drive efficiencies. Finally, since DsRed inheritance was lower than measured previously for sgRNA3 alone (97.4% male-drive and 93.9% for female-drive) (23), we tested gRNA6 alone with the same *U6* promoter-terminator. The Dm6 line produced the highest inheritance rates, with 90.8% for the male-drive and 75.2% for the female-drive. Male-drive was significantly higher than female drive (Fig 2, S3 Data, p<0.001) but lower than obtained with sgRNA3 alone.

### Target site disruption and formation of alleles that are potentially resistant to drive

The non-drive flies obtained from the homing assessment crosses were sequenced around the target sites to identify potential resistant allele formation due to Cas9-mediated cutting and error-prone repair (51, 52). For selected replicates, pooled non-drive individuals were deep-sequenced to measure indel frequencies. For the Dm126t line with X-linked Cas9, no modified alleles were detected with male-drive (n=30) whereas female-drive progeny showed 7.1%-7.9% modified reads around the targeted sites (S4 Data). Using autosomal Cas9, modification frequencies were 5.7% for male-drive (n=32) and 6.3% for female drive (n=31). Similarly, the Dm126t-term line showed low modification rates of 2.8% for male and female-drive. Higher modification rates were observed from female-drive of Ds36t (23.9%) and Ds36 (16.6%) (S4 Data).

For some crosses individual non-drive flies were Sanger sequenced and analyzed (see following section). Most of the modified alleles were large deletions including potentially non-functional (frameshift) alleles (Fig 3.), which suggests some expression and processing of mature sgRNAs from the multiplexed array. However, overall cutting efficiency was low, particularly for the lines with three sgRNAs.

**Fig 3.**
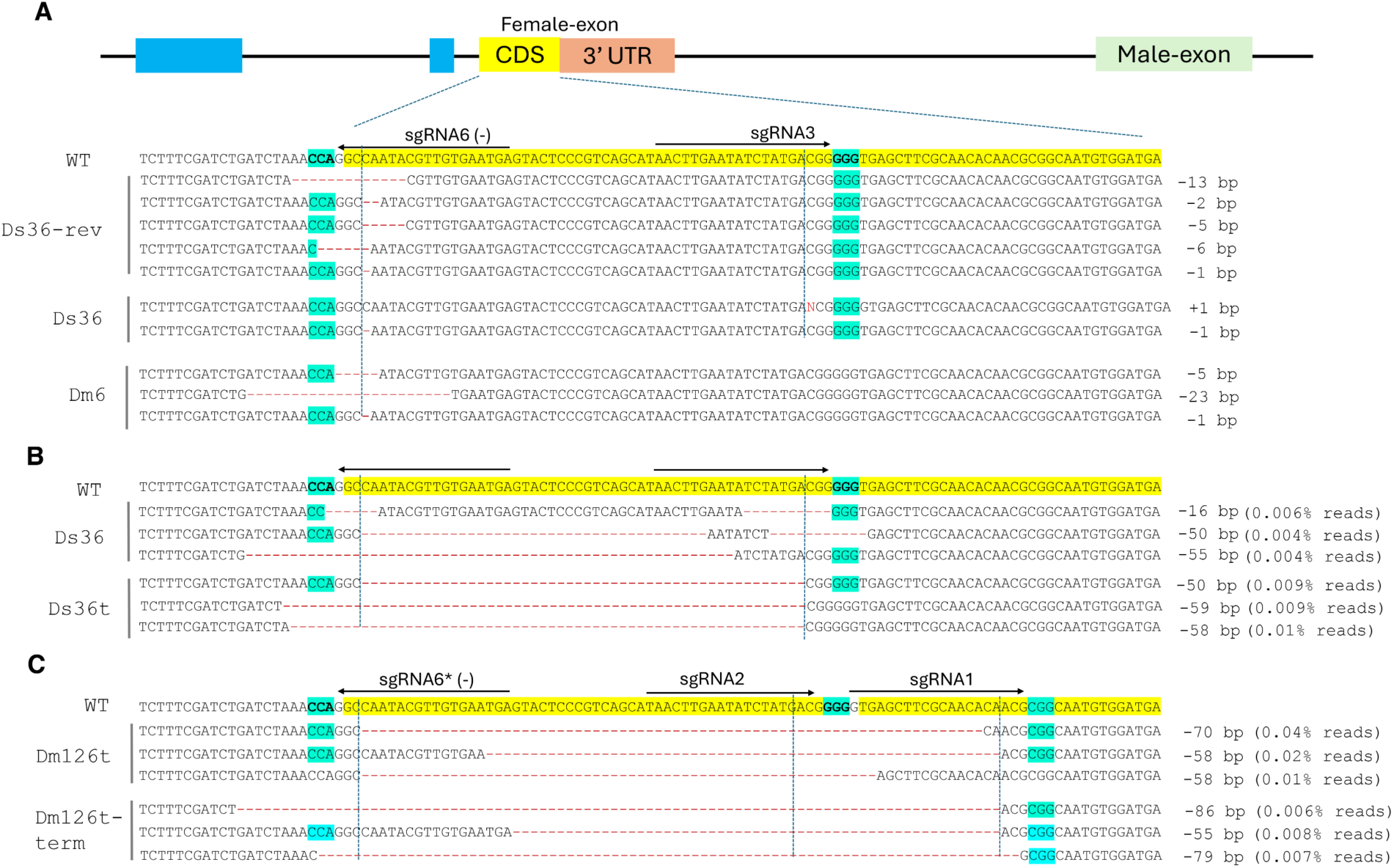
Characterization of resistance alleles at the *doublesex* target site. (A) Schematic of the *dsx* female-specific exon showing sgRNA target sites and representative potential resistance alleles identified in individual non-drive females by Sanger DNA sequencing. (B, C) Deep sequencing analysis of pooled non-drive progeny from constructs expressing two (B) or three (C) sgRNAs. Representative indel alleles are shown aligned to the wild-type sequence. Protospacer adjacent motif (PAM) sequences are highlighted in cyan, and vertical dotted lines indicate predicted Cas9 cleavage sites. Indel sizes and their relative frequencies are indicated.

### Dominant female sterility among non-drive progeny

In *D. melanogaster*, some female non-drive offspring from a *dsx* male drive showed dominant sterility (24). Interestingly, mutations that deleted the terminal AG of the splice acceptor site showed a stronger intersex phenotype than mutations with an intact AG (24). To assess similar effects in *D. suzukii*, fertility and genital morphology were examined in non-drive females from male-drive crosses set in bottles. For the Ds36-rev line, 6 of 51 non-drive females (11.7%) were sterile, with indels from 1-13 bp at the sgRNA6 target site (Fig. 3A, S5 Data). For the Dm6 line, 9 of 52 non-drive females (17.3%) were sterile, also associated with indels at the target site. In both strains, disruption of the splice acceptor motif produced stronger intersex phenotypes (S3 Fig.).

In contrast, for the Dm126t line with X-linked Cas9, all non-drive females (n=20) were fertile. The sequence analysis revealed wild type sequence for 16 individuals, but 4 females had a 1 bp (n=3) or 3 bp (n=1) insertion at the sgRNA1 target site. Since all mutant females were fertile, the mutations appear to be recessive or potentially not loss of function alleles (S5 Data). Similar results were obtained with autosomal Cas9 and for the Dm126t-term line with either Cas9 line (S5 Data). For Ds36 with X-linked Cas9, 5 out of 17 (29%) females were dominant sterile. Sequence analysis revealed a 1 bp deletion in 3 females around the sgRNA6 target site while one female showed 1 bp insertion at the sgRNA3 target site. In contrast, all females (n=16) for Ds36 with autosomal Cas9 were fertile as were the non-drive females (n=14) for the Ds36t line with X-linked Cas9. The numbers of fertile and sterile non-drive females obtained from male drive crosses are shown in S4 Fig.

We also assessed the fertility of non-drive female progeny from female-drive for some lines. For Ds36 with X-linked Cas9, 22 out of 36 (61%) were sterile and sequence analysis of 14 of the sterile females found mutations around the sgRNA6 (n=2) or sgRNA3 (n=2) target sites (S5 Data). For Ds36t (sgRNA6-tRNA-sgRNA3) 12 out of 21 (57%) females were sterile and sequence analysis revealed indels around target sites (S5 Data). The sterility among non-drive females of a female-drive could be due to maternal Cas9/gRNA activity (53, 54)

## Discussion

In this study, we evaluated a series of homing gene drive constructs targeting the female-specific exon of the *D. suzukii dsx* gene, comparing the effects of sgRNA number, expression architecture, and target-site selection on drive efficiency and resistance allele formation. We previously found very high homing rates in both sexes with a single sgRNA (sgRNA3) targeting a conserved region in the middle of CDS (Dm3, Fig 1.) (23). As previous studies have found it advantageous to target the conserved intron-exon boundary (18, 24), we evaluated a gRNA (sgRNA6) targeting this location alone or in combination with one or two sgRNAs targeting the CDS. When expressed alone, we found male drive efficiency was very high (90.8% inheritance), although less than obtained previously using sgRNA3 (97.4%) (23). Whereas we previously observed a very high drive efficiency in females with sgRNA3 (23), drive efficiency was significantly lower in females that expressed sgRNA6 alone. Similarly, in the strains that expressed both sgRNAs, drive efficiency was significantly higher in males than females. It is not clear why drive efficiency was higher in males than females but it may reflect differences in the preferential use of the HDR repair pathway in the male and female germlines. Previous work in *Drosophila* has shown that double-strand breaks in the germline and early embryos are frequently repaired by end-joining pathways rather than homology-directed repair (43, 55, 56), and that maternal deposition of Cas9 can shift cleavage to embryonic stages where homing of drive constructs does not occur. In *Culex quinquefasciatus*, homing gene drive frequencies were significantly higher in males than females, particularly if there was significant mismatch in the flanking regions (57). Since we did not observe a difference in drive efficiency in males and females with sgRNA3 (23), it would appear that it is cleavage at the sgRNA6 site at the intron-exon junction that is mostly responsible for the observed differences between the sexes.

Expression of a third sgRNAs lowered drive efficiency in males, with about 60% inheritance of the DsRed gene. Drive efficiency in females was similar and comparable to the two gRNA strains with about 55-60% inheritance of the marker gene. The decrease in male-drive could reflect limitations associated with increased guide number, such as competition among sgRNAs for Cas9 loading. However, the low drive rates could also be due to poor expression of sgRNAs due to limited tRNA processing. In particular, the leucine tRNA, which was only in the low-performing 3-gRNA constructs, may have had lower processing efficiency than the glycine tRNA, which was used in both the 3-gRNA and one of the 2-gRNA constructs. Consistent with this possibility, we observed a very low percentage of non-drive flies with modified reads, meaning that total cut rates were likely low. Recent mosquito studies have also indicated poor performance with ribozymes connecting gRNAs (49). Thus, for future drive designs that require expression of three of more sgRNA in *D. suzukii*, it would appear to be better to use separate polymerase III promoters for each sgRNA.

Individual gRNAs can also vary in activity, and it is possible that sgRNA2 (used in the 3-gRNA drives) had inferior activity compared to overlapping sgRNA3, which led to better male performance in the 2-gRNA drives. Indeed, sgRNA3 may be particularly effective, explaining the superior performance of Dm3 over the similar Dm6 construct. As the *DsU6d* promoter has has been reported to have higher activity than the *DsU6a* promoter (58, 59), this could also explain why the 2-gRNA construct with a tRNA did not have inferior performance to the other 2-gRNA systems.

Comparison of sgRNA expression architectures further showed that constructs expressing two sgRNAs from separate promoters performed similarly whether arranged back-to-back or separated within the construct, indicating that promoter orientation and spacing have minimal impact on homing efficiency. This provides flexibility in construct design and suggests that architectural considerations such as minimizing repetitive sequences or recombination risk may guide future designs without compromising performance.

A potential advantage of targeting the *dsx* intron-exon junction, was the generation of dominant female-sterile mutations among non-drive females as observed in *D. melanogaster* (24). Modeling indicated that the generation of these mutant alleles contributed to stronger and more rapid population suppression (60). While such mutations were found, the frequency of dominant sterility was lower than observed in *D. melanogaster*. This could suggest possible species-specific differences in *dsx* regulation or repair outcomes, though it could also have been caused by lower total cleavage rates in this study, leading to less simultaneous cleavage with different gRNAs. Indeed, most non-drive females were fertile and wild-type with a minority carrying small indels in the female exon. As in *D. melanogaster*, deletions that removed the splice acceptor site produced the strongest intersex phenotypes.

Homozygous males carrying the drive constructs were sterile, consistent with previous observations for *dsx*-targeting drives. Incorporation of a PEST degradation sequence, with or without a 3′ untranslated region, did not restore male fertility, indicating that this strategy is insufficient to mitigate fitness costs associated with *dsx* disruption. Recently, other strategies were tested in *D. melanogaster* such as inclusion of a *dsx* 3’UTR with TRA/TRA2 binding sites to promote female-specific splicing (60). However, none successfully rescued homozygous male fertility. The inability to produce fertile homozygous males has important implications for the expected performance of suppression drives in field populations. Because drive carriers would be effectively limited to hemizygous individuals, the rate of drive spread and population suppression may be reduced relative to systems in which homozygotes remain fully viable and fertile (60). This constraint could increase the release threshold required for population suppression in some self-limiting systems, and for self-sustaining systems, it could slow the rate of drive propagation and reduce the overall suppressive power of the system.

Together, our findings demonstrate that homing gene drive performance in *D. suzukii* is shaped by a combination of sgRNA number, expression architecture, target-site selection, and germline-specific repair processes. These results provide practical guidance for optimizing gene drive designs in *D. suzukii* and underscore the importance of testing multiple design strategies within the target species.

## Material and Methods

### Plasmids design and assembly

Plasmids HSDdsxU6:3Ds-1 (Dm126t) and HSDdsxU6:3Ds-3 (Dm126t-term) were designed and assembled in the Champer lab and were shipped to Scott lab spotted on filter papers. The plasmid containing spots were cut out individually using sharp clean blades and placed in 1.5 mL Eppendorf’s tubes containing 45 μL of elution buffer (Zymo, Cat#D3004-4-16) and incubated for 15 min at room temperature followed by centrifugation at 14,000g for 5 min at RT. Five μL of the eluate was used to transform NEB 10-β cells (NEB Cat#C3019H) following NEB transformation protocol. Further, mini preps (Zymo Cat#D4016) and midi preps (Zymo Cat#D4200-A, 4200-B) were performed to isolate the plasmids. Plasmid integrity was checked by restriction digestions and complete plasmid sequencing.

For assembly of HSDdsxU6:3Ds-1 and HSDdsxU6:3Ds-3 the *hsp83* promotor, *shu* 3’UTR and *dsx* left homology arm (1015 bp) and *dsx* right homology arm (1036 bp) were amplified from *D. suzukii* genomic DNA template by PCR. A GENEWIZ synthesize gRNA expression cassette, DsRed cassette and homology arms were cloned by Gibson assembly as an intermediate plasmid. The intermediate plasmid was digested with BsiW1 and BstZ171 and the myosin splicing acceptor sequence, PEST degradation tag and *shu* 3’UTR (Dm126t-term) were cloned by Gibson assembly.

Plasmid ‘pUC19-DsxHA-DsU6a-gRNA3-U6d-gRNA6-pUbDsRed’ (Ds36) was designed and developed to express two gRNAs under two *D. suzukii U6* promoters. An IDT synthesized gBlock fragment (1680 bp) containing the *dsx* left homology arm (1022 bp) and a myosin splice acceptor -hsp70 3’UTR sequence (23) was digested with BamHI and SacI was cloned into pUC19 vector digested with the same pair of enzymes to make the ‘pUC19-dsxLASA’ intermediate plasmid. The *dsx* right homology arm was amplified from *D. suzukii* genomic DNA template by PCR with primers dsxRF1 and dsxRR1 (S1 Table) that contain BamH1 and AgeI restriction sites. The Right arm was cloned into the pUC19-dsxLASA plasmid digested with BamHI and AgeI to generate the ‘pUC19-dsxLASaRA’ plasmid. The pUbDsRed maker was excised using AscI and AgeI from pUC57-DsxHDR-U6:3dsx-sgRNA3-pUbDsRed (23) and cloned into ‘pUC19-dsxLASaRA’ digested with AscI and AgeI to generate the ‘pUC19-dsxLASaRA-pUbDsRed’ construct. The sgRNA3 and sgRNA6 (S1 Table) sequences were sequentially cloned under the *DsU6a* and *DsU6d* promoters and their respective terminators following a Golden Gate cloning strategy used previously (23) by digestion of a synthesized plasmid ‘pUC57-DsU6a-DsU6d’ (Genscript) with BbsI and BsmBI to create ‘pUC57-DsU6a-gRNA3-DsU6d-gRNA6’ plasmid. This plasmid was digested with AscI, PspOMI and BsaI (this enzyme was used for better resolution of the desired fragment on gel) followed by gel purification of the ‘DsU6a-gRNA3-DsU6d-gRNA6’ fragment and cloned into the ‘pUC19-DsxLASaRa-DsRed’ plasmid digested with AscI and PspOMI to generate ‘pUC19-DsxHA-DsU6a-gRNA3-U6d-gRNA6-pUbDsRed’ plasmid.

Plasmid ‘pUC19-Dsx-tRNAg3,6-DsRed’ (Ds36t) was designed to express tRNA-linked two sgRNAs (the sgRNA6 and sgRNA3) under the *DsU6d* promoter. The IDT gBlock synthesized fragment ‘DsU6d-gRNA6-tRNA-gRNA3-U6d terminator’ (named dsxtRNA) was digested with PspOMI and AscI for the cloning steps. Plasmid pUC19-DsxHA-DsU6a-gRNA3-U6d-gRNA6-pUbDsRed was also digested with PspOMI and AscI and the desired fragment (pUC19-dsxHA-pUbDsRed) was gel purified and ligated with the dsxtRNA fragment to generate the ‘pUC19-Dsx-tRNAg3,6-DsRed’ plasmid (MS1).

Plasmid ‘pUC19-Dsx-gRNA3,6Flipped-DsRed’ (Ds36-rev) was designed to express two sgRNAs from spatially separated *U6* gene promoters. The plasmid Ds36 was digested with AgeI (sites flanking DsU6d-gRNA6-pUbDsRed) and re-ligated and screened for the ligation of a fragment in flipped orientation to generated ‘pUC19-Dsx-gRNA3,6 Flipped-DsRed’ (Ds36-rev) plasmid.

Plasmid ‘pUC19-dsxHADmU6:3-gRNA6-DsRed’ (Dm6) was designed to express only the sgRNA6 under the *D. melanogaster* U6:3 promoter and terminator (Addgene, #49410). The gRNA6 sequence was cloned into pCFD3 plasmid following Golden Gate cloning (BbsI). PCR was then used to obtain the ‘Dm6:3-gRNA6-terminator’ fragment. The primers, U6F1 and U6R1 (S1 Table), used contained PspOMI and AscI sites for subsequent cloning steps. Plasmid pUC19-DsxHA-DsU6a-gRNA3-U6d-gRNA6-pUbDsRed was also digested with PspOMI and AscI and desired fragment (pUC19-dsxHA-pUbDsRed) was gel purified and ligated with the Dm6:3-gRNA6-terminator’ fragment digested with same pair of enzymes to generate the ‘pUC19-dsx-DmU6:3-gRNA6-DsRed’ plasmid. Further, this plasmid wad digested with AgeI and AfIII to remove the right arm sequence (originally designed for homing using sgRNA3), the desired fragment gel purified followed by ligation to a right homology sequence (for the gRNA6 cut site) obtained by PCR amplification, using primers DsxAgeIF3 and DsxAfIIIR2 (S1 Table), of *D. suzukii* genomic DNA and digested with same pair of enzymes. The final Dm6 plasmid was identified by restriction enzyme analysis.

For all final plasmids, the integrity was determined by restriction digestion analysis and complete plasmid sequencing prior to embryo microinjection. The final plasmids were isolated from transformed *E. coli* (NEB 10 beta cells Cat#C3019H) using the Zymo Midi kit (Cat# D4200-A and D4200-B) followed by ethanol precipitation, washing with 70% ethanol and reconstitute the plasmids in nuclease free water to a final concentration of ∼1000 ng/μL. Also, prior to microinjection plasmids were filtered through 0.46 μm filter (MiliporeSigma, Cat# UFC30HV25).

### Embryo microinjection and establishment of homing gene drive strain

To make RNP complex for use with homing plasmids, synthetic crRNA:tracrRNA duplex (IDT synthesized) were used. For crRNA:tracrRNA, the dsxcrRNA (72 ng/μL) and tracrRNA (134 ng/μL) were mixed in nuclease free duplex buffer and incubated at 95 °C for 5 min and then allowed to cool to room temperature for duplex formation. The crRNA:tracrRNA duplex was mixed with Cas9 protein (IDT Cat#1081060) and incubated at 37 °C for 20-30 min. The final injection mix consisted of plasmid DNA at ∼500 ng/μL and Cas9 RNP complex at a final concentration of 500 ng/μL for Cas9. The injection mix was centrifuged at ∼20,000g for at least 10 min at 4 °C prior to injection. Pre-cellular embryos from the *D. suzukii* X-linked Cas9 (#27A1) strain were microinjected using quartz needles. The G_0_ adults flies that developed from injected embryos were individually crossed with wild type and the offspring screened for red fluorescence to identify positive transformants. Further crosses were made to remove the Cas9 carrying chromosome (green fluorescent), if present.

### Homing constructs knock-in confirmation

Genomic DNA was extracted from transgenic flies using either the Zymo DNA prep kit (Cat#D4069) or by a phenol chloroform extraction method (61). PCR amplification was performed using primer pairs (S1 Table) specific to the homing cassette and the flanking genomic regions for both sides of the insertion. The PCR products were column purified (Zymo Kit Cat#D4014) and sequence determined using Sanger DNA sequencing. PCR, using primers (S1 Table) for sgRNA cassette regions, and Sanger sequencing were also performed to confirm the integrity of the sgRNA transgenes upon successful knock-in of the homing construct.

### Fly rearing and homing assessment crosses

All established homing strains were raised on cornmeal-yeast-agar diet at room temperature (20-22°C). *D. suzukii* diet was prepared as follows: 70 g agar, 300 g sucrose, 400 g cornmeal, and 200 g dry yeast were added to 8 L of water and mixed. The mixture was heated to boil, 250 mL molasses was then added and the heat turned off. After 20 min, 24 mL of 10% Moldex (Cat#Sigma-Aldrich H5501-500G) (in 95% ethanol) was added and mixed well. The diet was poured into vials and bottles, ready to for use the next day. To measure homing drive inheritance, previously characterized X-lined Dsnos-NLS-Cas9-NLS (#27A1) and autosomal (2^nd^-chromosome) Dsnos-NLS-Cas9-NLS (#25A2) virgin females (23) were crossed with males of homing strains to obtain trans-heterozygous (carrying Cas9 and gRNA) F_1_ progenies. The trans-heterozygous male and virgin females, called “male-drive” and “female-drive” respectively, were further crossed with wild type flies of the opposite sex. The F_2_ progeny were scored for the inheritance of the red-fluorescent protein marker using an M205FA microscope (Leica Microsystem, Buffalo Grove, IL) with DsRed filter (ex 545/25, em 595/50 nm). The F_2_ progeny that did not show red fluorescence were called “non-drive flies” and were quick frozen and then stored at -80 °C for future molecular analysis. All homing assessment crosses were performed in triplicate. The gene drive experiments were performed in an ACL2 containment facility.

For some of the crosses, the fertility status of the female “non-drive flies” were also tested before freezing them. Individual females (3-5 days old) were crossed with wild type males (n=3) and were kept in food-vials for at least a week. The food-vials were monitored for any offspring for at least two weeks.

### Screening for resistance alleles

Genomic DNA was isolated from the non-drive flies using phenol-chloroform extraction method described previously (61). A region of the *dsx* gene was amplified using Q5 polymerase 2X master mix (NEB#0429S) and the primers dsxcutF1/dsxcutR1 or dsxcutF2/dsxcutR1 (S1 Table) and the purified PCR product sequenced using Illumina DNA-sequencing. The deep sequencing data were analyzed using CRISPResso2 software (62). Pair-end reads with a minimum average read quality 30 were analyzed for indels on the target sites. DNA was also extracted from individual flies by homogenizing in 50 μL squash buffer (10 mM Tris pH8.2, 1 mM EDTA pH8, 25 mM NaCl and 200 μg/mL proteinase K) followed by incubation at 37 °C for 45 min and 95 °C for 2 min. PCR was performed as above with 1 μL of genomic DNA and the product column purified. DNA sequences were determined using Sanger sequencing and analyzed using Synthego-ICE tool (Edit-Bio) (63).

### Statistical analysis

Drive inheritance was calculated as the proportion of DsRed-positive F2 progeny among all scored F2 progeny. As the data arise in the form of standard binomial data we fitted a generalized linear mixed model (GLMM) using R (4.5.2) (S3 Data). The lme4 (64) and emmeans (65) packages were used for model fitting, parameter estimation, and statistical inference. The response distribution of the GLMM was specified to be binomial, and the logit link function was used. The fixed effects included were the effect of treatment (drive line/cas9 combo), sex, and the interaction of treatment and sex. Experimental runs were conducted in batches, therefore, a random effect associated with the batch was also included. The target distribution was the conditional distribution of the observations given the batch effects. 15 quadrature points were used for Gaussian quadrature. The final model did not show any indication of problematic fitting.

## Acknowledgements

We thank Aki Yamamoto for assistance with setting *Drosophila suzukii* crosses, James Clothier for statistical analysis of the inheritance data, Esther Belikoff for guidance on ACL2 containment and our colleagues in the Scott lab for helpful discussions.

This work was supported by Biotechnology Risk Assessment Research (BRAG) program grants 2021-33522-35341 and 2024-33522-42694 from the USDA National Institute of Food and Agriculture to MJS.

## Declaration of generative AI and AI-assisted technologies in the writing process

During the preparation of this work, the corresponding author (MJS) used ChatGPT (5.5) in order to improve the readability of the manuscript. After using this tool/service, the author reviewed and edited the content as needed and take full responsibility for the content of the published article.

