## Supplementary material for "Target site and guide RNA multiplexing architecture shape homing gene drive efficiency in *Drosophila suzukii*": S1 Fig

### Supplementary Figures

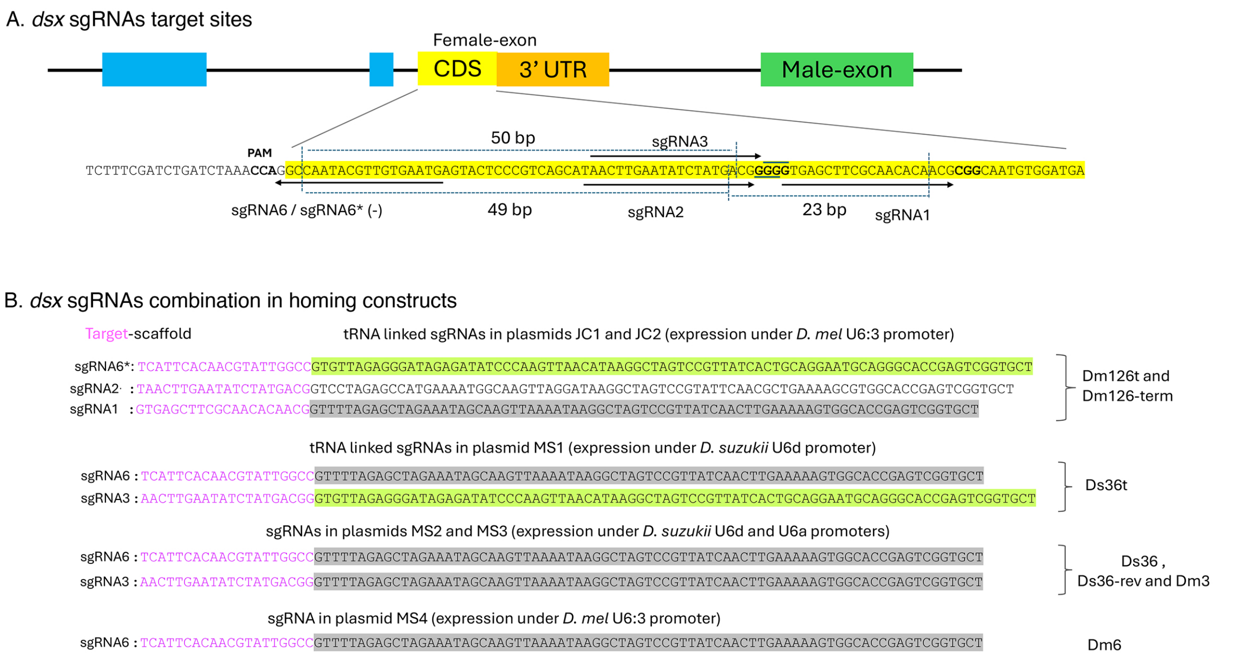


**S1 Fig. sgRNA target sites and organization of multiplexed guide RNAs in homing constructs targeting *doublesex*.** (A) Schematic of the *doublesex* (*dsx*) locus showing sgRNA target sites within the female-specific exon with coding sequence highlighted in yellow. Distances between adjacent target sites are indicated. Protospacer adjacent motif (PAM) sequences are highlighted. (B) Organization of sgRNAs within homing constructs. In Dm126t and Dm126t-term, three sgRNAs are expressed as a single transcript under the *D. melanogaster* U6:3 promoter and separated by tRNA sequences. Ds36t, Ds36 and Ds36-rev express sgRNA3 and 6 using a tRNA (Ds36t), or independent *D. suzukii* U6 promoters (Ds36, Ds36-rev). In Dm6, a single sgRNA is expressed under the *D. melanogaster* U6:3 promoter. Distinct sgRNA scaffold sequences are indicated, and homing strains derived from each construct are shown on the right.

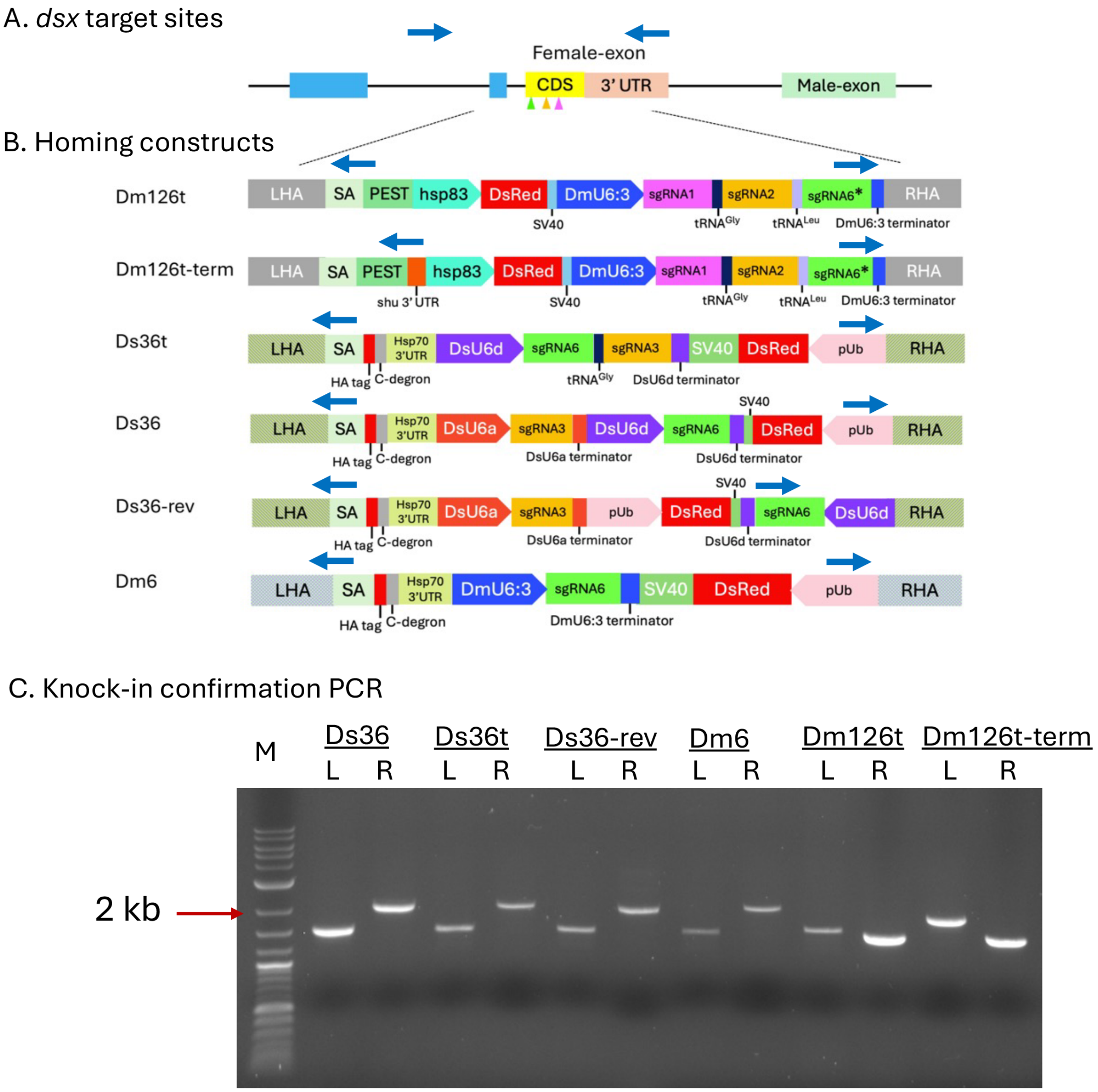


**S2 Fig. PCR confirmation of homing construct integration at the *doublesex* locus.** (A–B) Schematic representation of the *dsx* female-specific exon and homing construct integration sites (not to scale). Blue arrows indicate primer positions used for diagnostic PCR across the left (L) and right (R) homology arms. (C) Representative agarose gel showing PCR products confirming correct insertion of homing constructs at the *dsx* locus. M, 1 kb Plus DNA ladder (NEB, N0550S). For each construct, amplification was performed across the left (L) and right (R) junctions of the knock-in site.

Expected amplicon sizes were as follows:

Left junction—Ds36, Ds36t, Ds36-rev, and Dm6: 1504 bp; Dm126t: 1567 bp; Dm126t-term: 1845 bp.

Right junction—Ds36, Ds36t: 2041 bp; Ds36-rev: 2000 bp; Dm6: 2093 bp; Dm126t and Dm126t-term: 1438 bp.

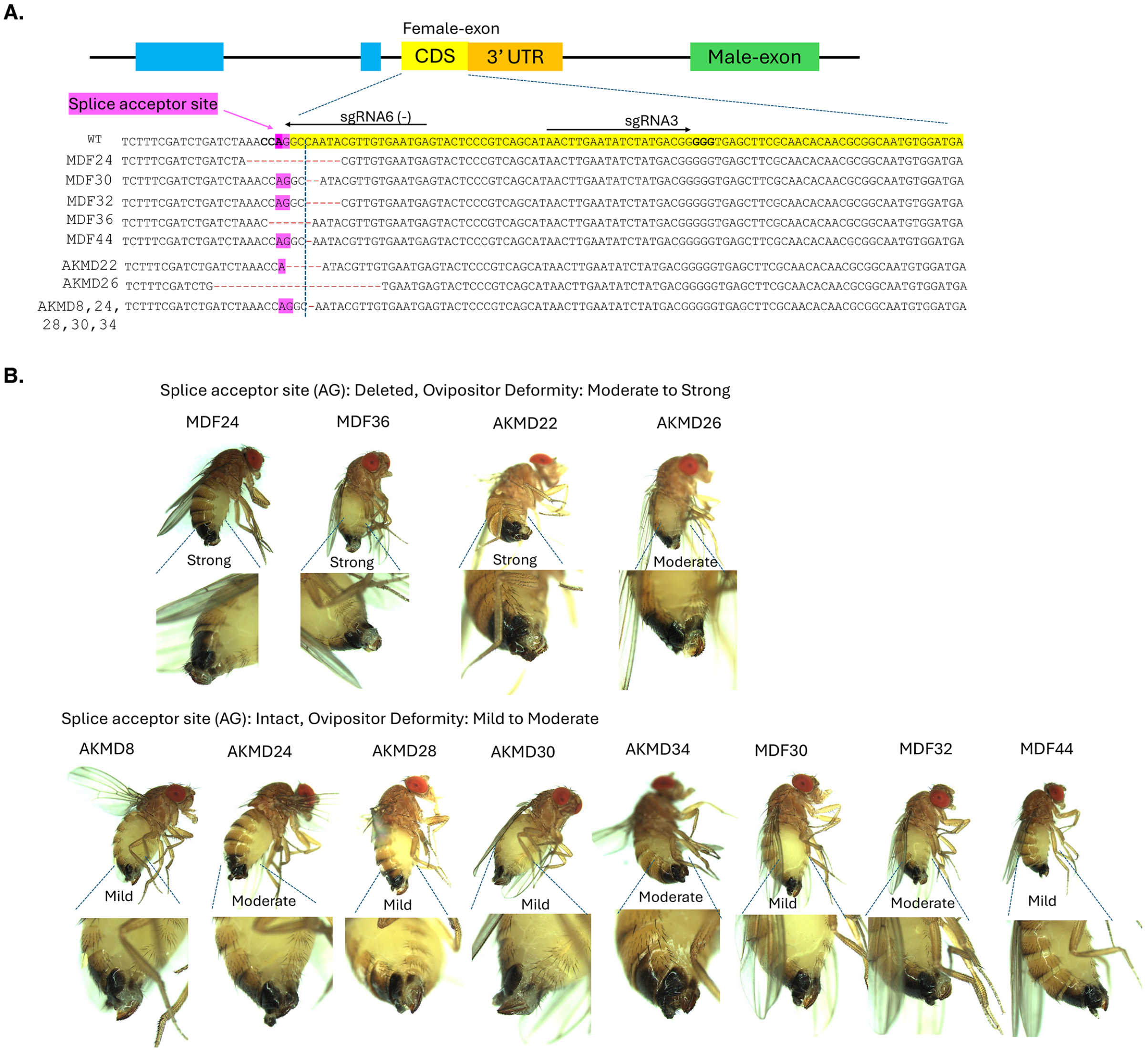


**S3 Fig. Dominant female-sterile resistance alleles and associated ovipositor phenotypes.** (A) Schematic of the *doublesex* (*dsx*) female-specific exon showing sgRNA target sites and representative indel alleles identified in dominant sterile non-drive females. The splice acceptor site (AG) of the female intron s highlighted. (B) Representative ovipositor morphology of dominant sterile females carrying resistance alleles. Deletion of the splice acceptor site (AG) is associated with moderate to severe ovipositor deformities, whereas mutations that retain the AG sequence result in mild to moderate phenotypes. Individual samples are labeled by line and identifier (e.g., MDF for Ds36-rev male-drive; AKMD for Dm6 male-drive).

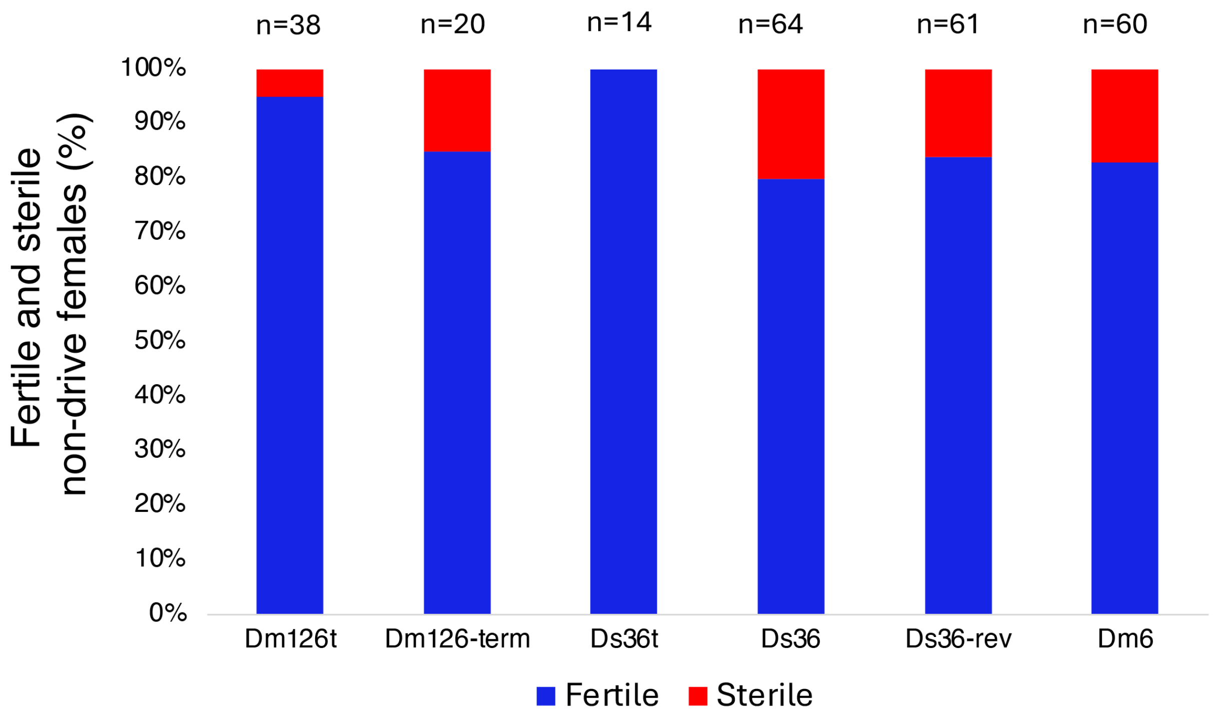


**S4 Fig. Fertility status of non-drive females obtained from male-drive crosses.** Numbers (n) represent the total counts of non-drive females pooled from all male-drive crosses using autosomal and X-linked Cas9 lines.

**S1 Table. Oligonucleotides/Primers used in this study**

| Name | Sequence (5'-3') | Use |
|---|---|---|
| dsxRF1 | ATCTAATGTAACCGGTCGGGGGTGAGCTTCGCAACACA ACGCGGCA | PCR out *dsx* right homology arm |
| dsxRR1 | ACTATAGTATTGGATCCTTGTGCACCATCTCACAACAGT TTGATAC | |
| DsxAgeIF3 | ATATTCACCGGTTATCAATACGTTGTGAATGAGTACTCC CGTC | PCR out *dsx* right arm of plasmid Ds36-Rev |
| DsxAflIIR 2 | AAGTACTTAAGTAAAGTTAAAATAACATGAATGACTGA CGTG | |
| pCFD3_g6 SS | GTCGTCATTCACAACGTATTGGCC | Cloning sgRNA6 |

| | | |
|---|---|---|
| pCFD3_g6 AS | AAACGGCCAATACGTTGTGAATGA | under DmU6:3 |
| DsU6d_g6 SS | ATCGTCATTCACAACGTATTGGCC | Cloning gRNA6 under DsU6d, used with pCFD3_g6 AS |
| DsU6a_g3 SS | TTCGAACTTGAATATCTATGACGG | Cloning sgRNA3 under DsU6a |
| DsU6a_g3 AS | AAACCCGTCATAGATATTCAAGTT | |
| DsxcutF1 | GGGTAAGTGAACTGATCATTGAAAC | PCR out target region |
| DsxCutR1 | GAAGATAATCCAAGTTGCGCTTTAG | |
| DsxcutF2 | GTATGTGATATTGAAGGATGCAGAC | |
| DsxcutR2 | CGCTTTAGCTTACCGATAAATTCG | |
| U6F1 | TGAATAAGGGGCCCTGCCATACCATTTAGCCGATCA | PCR out DmU6:3 gRNA6 region for cloning |
| U6R1 | ATAGATAAGGCGCGCCGAGCACAATTGTCTAGAATG | |
| DsxHDF7 | GCGGGCACAATATTAATACAAA | Ds36, Ds36t, Ds36-rev and Dm6 left site knock-in confirmation |
| DsxSALA_R1 | GTAGGCTGCGGAAGAGAGATAAAT | |
| DsxHDR_F3 | AGTGTGATGTGTTGAGCGTTG | Ds36, Ds36t, Dm6 right site knock-in confirmation PCR and with DsU6d_SeqR for Ds36-rev |
| DsxHDR_R3 | AATCCCTGAAACCTGCCTCTA | |

| | | |
|---|---|---|
| | | right site PCR |
| RR1_U6aF1 | TTTGGTTAGGGACCAAAGAAAGT | DsU6a-gRNA3 region PCR |
| RR1_U6dF2 | TCGTAGACCCTTTCGATCTCATA | |
| DsU6d_SeqR | TGTTATTCACCTCAGGTTGGCAA | DsU6d-gRNA6 region PCR |
| DsU6d_SeqF | GCCAGTGCTGAATTCTGTAACTG | |
| RR1_U6aR2 | GCAATAGCATCACAAATTTCACA | |
| RR1_U6dR1<br>DsU6d_SeqR | ACAGCACAATCAACTCAAGAACA<br>TGTTATTCACCTCAGGTTGGCAA | |
| JLR1 | TCACTTGGTATCATCCAGATCC | Dm126t left site PCR with DsxHDF7 |
| JRF2 | TATAGACAATGGTTTTCCGTTGA | With DsxHDR_R3, Dm126t right site PCR |
| JCF1 | CGCCAGTGCTCACTACTTTTTAT | Dm126t and Dm126t-term gRNAs region PCR and sequencing |
| JCR1 | TATGTACGTCAACGGAAAACCAT | |
| JCF2 | ACCTGTGATTGCTCCTACTCAAA | |
| JCR2 | ATTCCTGCAGTGATAACGGACTA | |
| JC2R1 | TGTTAAGTAGTTGCAGGTAACCAT | Dm126t-term left site PCR |

| | | |
|---|---|---|
| | | with DsxHDF7 |
| JRF2 | TATAGACAATGGTTTTCCGTTGA | Dm126t-term right site PCR with DsxHDR_R3 |
| AKUF | TAGAATGAAACGCCACCTACTCA | Dm6 gRNA region PCR |
| AKUR2 | GCATACGCATTAAGCGAACATTA | |
| FMU6aF1 | TCGTAGACCCTTTCGATCTCATA | Ds36-rev gRNA region PCR |
| FMU6aR1 | GCAAGTCGAACCAAGTATTCAAG | |
| FMU6dF2 | TGTTCCCTTTAGTGAGGGTTAAT | |
| FM6dR2 | ACAATACCAGACCCCTTCTGTTT | |
| Dsdsx_LAF | GGTCGTCGTCGTCAAATTTA | PCR out *dsx* left homology arm for Dm126t and Dm126t-term |
| Dsdsx_LAR | GCCTGGTTTAGATCAGATCGA | |
| Dsdsx_RAF | ACGCGGCAATGTGGATGATA | PCR out *dsx* right homology arm for Dm126t and Dm126t-term |
| Dsdsx_RAR | TCCGTTGGGCTAAATCTTACT | |
| Dsshu_F | ATTTACTAAACAAACCAGTATTACA | PCR out *D. suzukii shu* promotor for Dm126t and Dm126t-term |
| Dsshu_R | CGGAATAACATATTCTCTTTCC | |
| Hsp83_F | CCATTTGTTCGGGGGCA | PCR out *D. suzukii* |

| | | |
|---|---|---|
| Hsp83_R | CTAGCAAGGGATTAGATCAATAGGA | *hsp83* promotor for Dm126t and Dm126t-term |
